# SAM68-regulated ALE selection of Pcdh15 maintains proper synapse development and function

**DOI:** 10.1101/2023.04.04.535307

**Authors:** Mohamed Darwish, Masatoshi Ito, Akinori Takase, Noriko Ayukawa, Satoko Suzuki, Masami Tanaka, Yoko Iijima, Takatoshi Iijima

## Abstract

Thousands of mammalian genes encode alternatively spliced isoforms in their 3’ untranslated region (3’UTR). Alternative 3’UTR diversity may contribute to several neurological processes in developing and adult brains. SAM68 is the key splicing regulator for the diversity of neuronal 3’UTR isoforms through alternative last exon (ALE) selection. However, the mechanisms underlying the control of splicing at the 3’ end and its function in the nervous system remain unclear. Here, we show that neuronal SAM68-dependent ALE splicing is regulated depending on its target transcripts. For example, the selection of the ALE of *protocadherin-15* (*Pcdh15*), a gene implicated in Usher syndrome and several neuropsychiatric disorders, is largely dependent on the expression of SAM68, partially regulated via the CaMK pathway, but independent of the U1 small nuclear ribonucleoprotein. We found that the aberrant ALEs of *Pcdh15* caused membrane-to-soluble isoform conversion of the produced protein and disrupted its localization into excitatory and inhibitory synapses. In addition, the neuronal expression of the soluble form of PCDH15 (sPCDH15) preferentially affected the number of inhibitory synapses. sPCDH15 further reduced neuroligin-2-induced inhibitory, but not excitatory, synapses in artificial synapse formation assays. Our findings provide insights into the role of alternative 3’UTR isoform selections in synapse development.

## Introduction

Thousands of mammalian genes encode alternatively spliced isoforms in their 3’ untranslated region (3’UTR). Alternative 3’UTR isoforms are generated through alternative last exon (ALE) splicing and alternative polyadenylation (APA) (1,2). Most human genes have multiple ALE and polyadenylation sites (PASs), which enable the expression of distinct 3’UTR isoforms in a tissue- or cell-type-dependent manner (3). The differences in the 3’UTR through ALE splicing and APA may be linked to the establishment of cell identities (4) and are accompanied by cell development (5,6). Thus, ALE splicing and APA are dynamically regulated in a spatiotemporal manner and are highly implicated in human diseases including hematological, immunological, and neurological diseases and cancer (7). Transcriptomic diversity through the spatiotemporal and dynamic regulation of the ALE and APA contributes to biological processes such as cell differentiation and identification. Information in the 3’UTR regulates mRNA targeting, translational efficiency, and stability (8). In neurons, specific mRNAs are localized near synapses, where localization is regulated by ALE splicing and APA (6). This regulation of local translation allows neurons to deal rapidly with synaptic functions and underlies important cellular processes such as memory dendrite formation, synapse formation, and axon guidance (9). ALE selection can change protein isoforms by truncating the coding sequence and/or providing alternative carboxy-terminal extensions of the protein, which dynamically change its functions (10). The usage of the ALE may underlie functional diversity at the transcriptomic and proteomic levels. However, the tissue- and/or cell-type specific mechanisms underlying the control of ALE splicing and APA in the nervous system and their subsequent functions remain unclear.

Src associated in mitosis (SAM68; 68 kDa) is a KH domain RNA-binding protein that belongs to the signal transduction and activation of RNA family. SAM68 is a tissue-specific key regulator of the diversity of neuronal 3’UTR isoforms through ALE splicing and APA as the knockout of *Sam68* preferentially causes premature termination at internal PASs (10–12). SAM68 synergistically acts with U1 small nuclear ribonucleoprotein (U1 snRNP) on pre-mRNAs to prevent improper termination in specific tissues (11,13) by preventing cleavage and polyadenylation at cryptic PASs through a process called “telescripting” (14). The interaction between SAM68 and U1 snRNP ensures proper 3’ processing during germ cell differentiation (11). Thus, the mechanisms by which SAM68 regulates ALE and APA and its subsequent effects on neuronal function are beginning to be uncovered.

We previously found that SAM68 regulates the selection of ALEs in several genes, including *protocadherin-15* (*Pcdh15*) (10). PCDH15 is a non-clustered protocadherin that plays an essential role in the maintenance of retinal and cochlear function (15). The gene is responsible for Usher syndrome, which presents with hearing loss following retinal degeneration (16,17). Mutations in the *Pcdh15* gene, including copy number variations (CNVs) and single nucleotide polymorphisms (SNPs), are also implicated in autism, bipolar disorder, and schizophrenia (18–21). However, it is unknown how the aberrant SAM68 splicing of *Pcdh15* can cause brain deficits. Here, we reveal that the neuron-specific ALE splicing of *Pcdh15* by SAM68 is regulated in a distinct manner that is different from that of other SAM68- regulated transcripts. In addition, the aberrant ALE of *Pcdh15* perturbs synapse formation and function through the disruption of proper protein localization into excitatory and inhibitory synapses.

## Results

### Dose-dependent ALE isoform selection of target transcripts by SAM68

We previously found that the deletion of SAM68 results in a long-to-short isoform switch of several neuronal targets through alterations in ALE selection (10). RNA sequencing (RNA-seq) and exon array analyses of *Sam68*/*Slm1* ^KO^ brains revealed that four SAM68-targeted transcripts, including interleukin 1-receptor accessory protein (*Il1rap*), ceruloplasmin (*Cp*), *Pcdh15*, and leucine-rich repeated coiled-coil protein 1 (*Lrrcc1*), are exclusively shortened at the 3’ end through altered ALE selection (Figures 1A and S1). This effect was specifically due to the deletion of SAM68 but not SLM1 (10). We further checked the dose dependency of the SAM68- targeted transcript*s* on SAM68 between various tissues and found that the amounts of the short forms of these transcripts were inversely correlated to the expression of *Sam68* (Figure 1B). The expression of the short forms of these transcripts did not appear to be dependent on *Slm1* (Figure 1C). The ALE isoform selection of the target transcripts of SAM68 is therefore dose- dependent.

**Figure 1:**
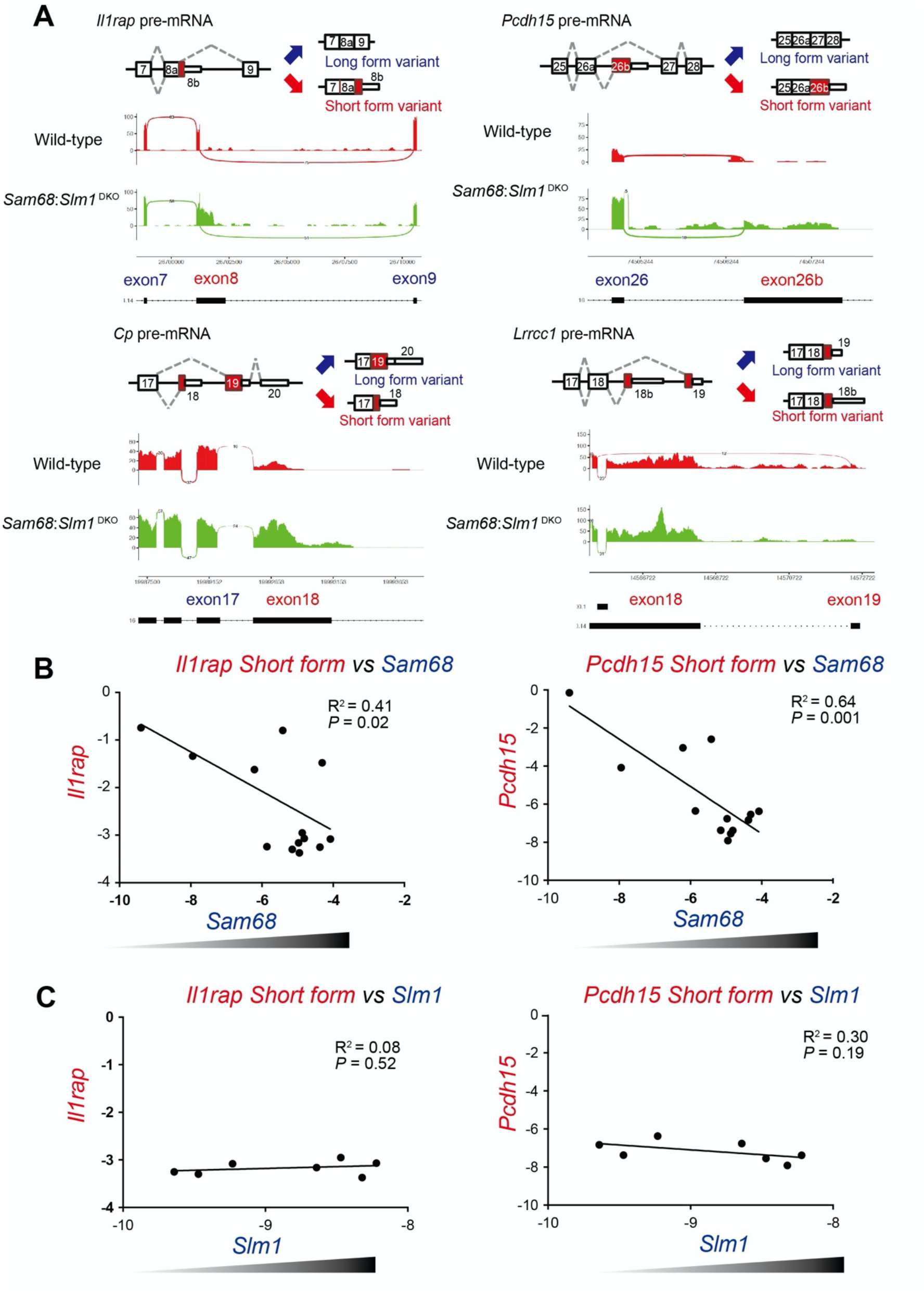
SAM68-dependent ALE selection of several transcripts in the central nervous system. (A) Aberrant 3’UTR exon selection of the representative genes in Sam68/Slm1^DKO^ brains. The upper panels show a schematic illustration of the selection of the ALE in four representative genes targeted by SAM68, including *Il1rap* (exon 8), *Cp* (exon 18), *Pcdh15* (exon 26b), and *Lrrcc1* (exon 18b). Lower panels show sashimi plots of the ALE events in the relevant genes. Each plot includes cassette exons and intron retentions. The red plots represent the wild-type mice, and the green plots represent *Sam68*/*Slm1* ^DKO^. The X-axes show genomic loci, and the Y-axes indicate transcription intensity. A “sashimi-like” region in each plot indicates an exonic region, and the blank regions between them indicate intronic regions. The numbers on the bridges crossing exons indicate junction reads. The raw data [the DDBJ database (6781), with accession number of DRA6781, with bioproject accession number of PRJDB6781] was obtained in the previous study (10). The RNA-seq was based on the UCSC genome browser Mouse NCBI37/mm10 assembly. (B, C) Correlation between *Sam68* (B) or *Slm1* (C) transcript levels and the production of *Il1rap* and *Pcdh15* short forms in various brain regions and tissues. ΔΔCT values of short form transcripts were quantified by RT-qPCR analyses and compared to those of *Sam68* or *Slm1*. The correlation coefficient (R^2^) and significance level between *Sam68* or *Slm1* and the SF transcripts are shown in the scatter plot.

### Different neuronal activity dependency of ALE isoform selections of SAM68-targeted transcripts

We previously showed that SAM68 regulates neuronal activity-dependent alternative splicing (AS) (22); therefore, we assessed whether SAM68-regulated ALE splicing can be affected by neuronal activity. We analyzed the effects of neuronal activity on the ALE selection of SAM68 targets *in vitro* by treating differentiated cortical neuron cultures with bicuculine (Bic) or tetrodotoxin (TTX) to alter spontaneous neuronal activity (Figure 2A). There were significant increases in the short forms of *Pcdh15* and *Lrrcc1* and inversed decreases in the long forms of *Lrrcc1* with TTX. The long and short forms of *Il1rap* and *Cp* were not significantly altered by either treatment (Figure 2B). The ALE of various SAM68-targeted transcripts is therefore influenced by spontaneous neuronal activity.

**Figure 2:**
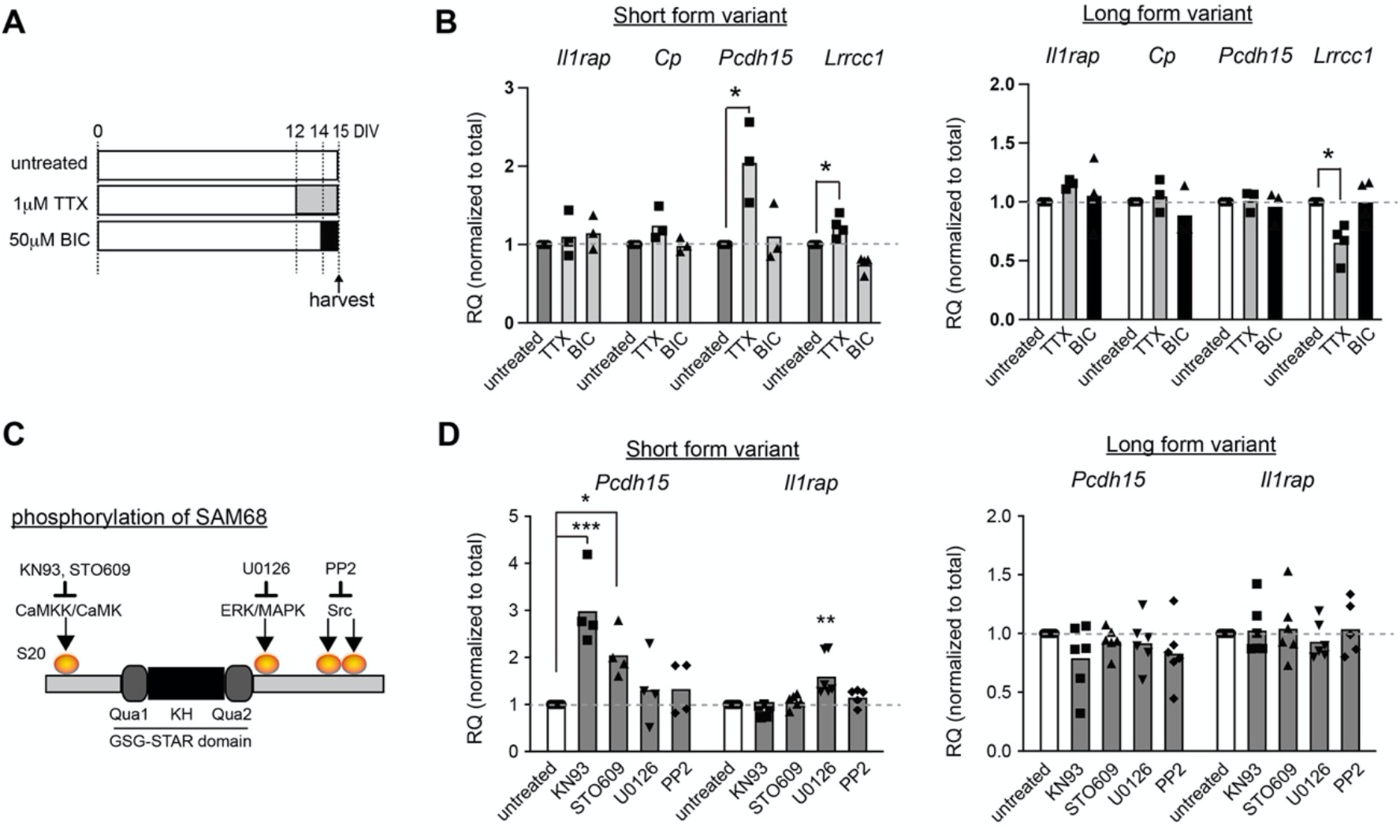
Distinct regulation of SAM68-regulated ALE splicing by neuronal activity and cellular signals. (A) Schematic overview of the pharmacological treatment protocol. Cultured cortical neurons were treated with bicuculine (BIC) (50 μM for 1 day) and Tetrodotoxin (TTX) (1 μM for 3 days) before harvesting on DIV15. (B) Relative expression of the short- and long-form variants of *Il1rap*, *Cp*, *Pcdh15,* and *Lrrcc1* shown in Fig 1A. n = 4–6 experiments per condition. Tukey’s multiple comparisons test was followed by a one-way ANOVA. (C) Schematic illustration showing SAM68 phosphorylation by several cellular signals/kinases and their inhibitors (KN93, STO609, U0126, or PP2). (D) Relative expression of the short- and long-form variants of *Pcdh15* and *Il1rap* treated with kinase inhibitors are shown in (C). Cultured cortical neurons were treated with inhibitors on DIV14 and harvested on DIV15. n = 4–6 experiments per condition. A one-way ANOVA was followed by Tukey’s multiple comparisons test.

Since SAM68 contains several phosphorylated sites that modulate its splicing activity, we analyzed whether blocking their activities would affect the ALEs of the target transcripts of SAM68 (23) (24) (Figure 2C). The short form of *Pcdh15* was significantly increased by CaMK inhibitors, KN93, and STO609, but did not significantly change by the actions of ERK/MAPK inhibitors, U0126, or the Src inhibitor, PP2 (Figure 2D). The ALE of *Il1rap* did not significantly change by the actions of any of these inhibitors. The ALE of *Pcdh15* may therefore be controlled by CaMK activity and the blockade of spontaneous activity (Figure 2B).

### Different usage of U1 snRNP on ALE isoform selections in SAM68-targeted transcripts

We next assessed whether the splicing of SAM68-targeted transcripts is regulated by interactions with U1 snRNP (11) (13). U1 snRNP binds to multiple intronic sites near cryptic PASs and prevents premature termination on a genome-wide scale (14) (25). We found U1 motif-like sequences at genomic loci around the cryptic PASs of *Il1rap*, *Lrrcc1,* and *Cp* (Figures 3A, 3B, and S2A), but not around those at exon 26b of *Pcdh15*. We then tested the alterations in the ALE selection of *Il1rap* and *Pcdh15* upon the depletion of U1 snRNP. U1 antisense morpholino oligonucleotides (U1 AMOs) and control AMOs were electroporated at different concentrations into primary cortical cultures. ALE selection was then assessed by quantitative PCR (qPCR) at day in vitro (DIV)-4 (Figure 3C). There was a significant concentration-dependent increase in the short form variant of *Il1rap* and reductions in its long form variants in U1 AMO-treated neurons. The total amount of *Il1rap* did not change, which indicated that the observed alteration was due to a shift in long-to-short isoform conversion (Figure 3C). The *Pcdh15* variants were not significantly altered (Figure 3D). We also checked the numbers of neurons and synapses in the absence and presence (20 µM) of U1 AMOs. There was a significant difference in the number of synapses but no difference in that of neurons (Figure S2B and S2C), indicating that U1 AMO electroporation is not lethal to neuron cultures under these conditions. Thus, the usage of U1 snRNP is different between targeted transcripts, which is consistent with the aforementioned findings on activity dependency. Together with the data in Figure 2, the results suggest that SAM68-dependent ALE selection is regulated in a distinct manner that is dependent on the target transcript.

**Figure 3:**
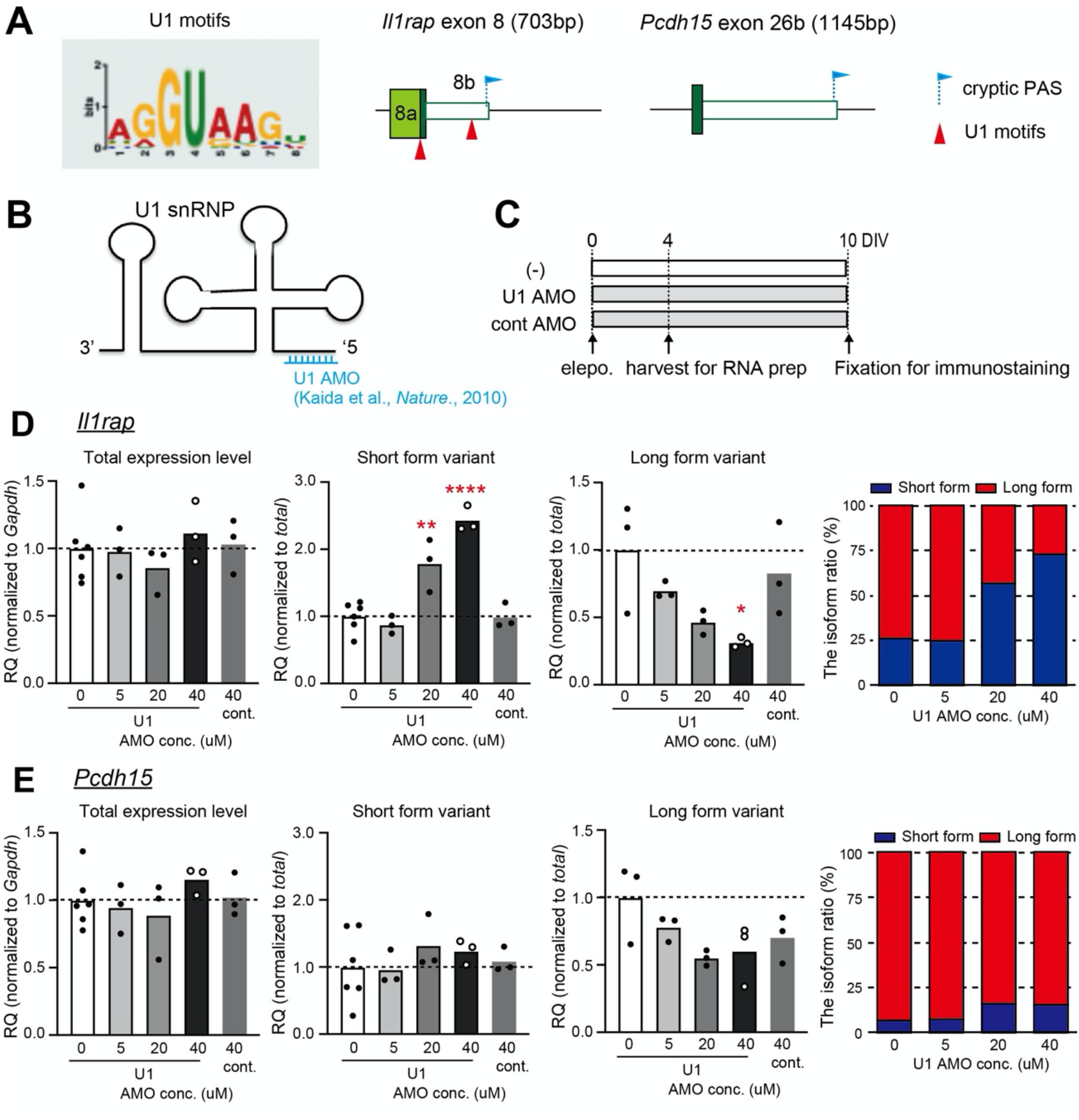
Distinct usage of U1 snRNP on ALE isoform selections of SAM68- targeted transcripts. (A) Logo representing the U1-binding consensus sequence (U1 motif) scanned for motif analysis enrichment (left). Schematic illustration showing the candidate U1 motif positions around ALE sequences in *Il1rap* (middle) and *Pcdh15* (right). U1 motifs and the cryptic polyadenylation sites (PAS) of each exon are labeled in arrowheads and flags, respectively. (B) Schematic representation of U1 snRNP. The antisense oligonucleotide binding site at the 5’ end is shown in blue (U1 AMO). (C) Schematic overview showing the protocol used for the U1 AMO-treated cortical neuronal culture. U1 or control AMOs were electroporated into plated cells at DIV0. The cells were harvested at DIV4 for RT-qPCR or further maintained until DIV10 for immunostaining. (D, E) Relative expression of the total mRNA, long-form (LF) and short-form (SF) variants, and abundance ratio of the SF (red) to LF (blue) between different concentrations of U1 AMOs by RT-qPCR. The RQ value of the total transcripts was normalized to *Gapdh*, and the RQ value of each alternative isoform was normalized to the total transcripts. n = 3–6 experiments per condition. A one-way ANOVA was followed by Tukey’s multiple comparisons test. (D) *Il1rap.* Total expression: one-way ANOVA [F (2, 9) = 0.89, P = 0.44], LF variant: one-way ANOVA [F (4, 10) = 3.75, P = 0.04]. SF variant: one-way ANOVA [F (4, 13) = 26.18, P < 0.0001]. (E) *Pcdh15* total expression: one-way ANOVA [F (4, 10) = 2.51, P = 0.11], LF variant: one-way ANOVA [F (4, 10) = 2.51, P = 0.11]. SF variant: one-way ANOVA [F (4, 13) = 26.18, P < 0.0001].

### Aberrant ALE selection of Pcdh15 alters its subcellular localization in neurons

We analyzed altered SAM68-dependent ALE splicing at the protein level. Previously, we found that the altered ALE usage of *Il1rap* results in the conversion of a membrane-bound to secreted type in *Sam68* ^KO^ brains, which results in alterations in *Il1rap* neuronal functions (10). We therefore assessed whether the aberrant ALE selection of *Pcdh15* affects its cellular localization. Similar to *Il1rap*, an insertion of exon 26b led to the conversion of PCDH15 from a membrane-bound form (mPCDH15) to secreted soluble form (sPCDH15) (Figure 4A).

**Figure 4:**
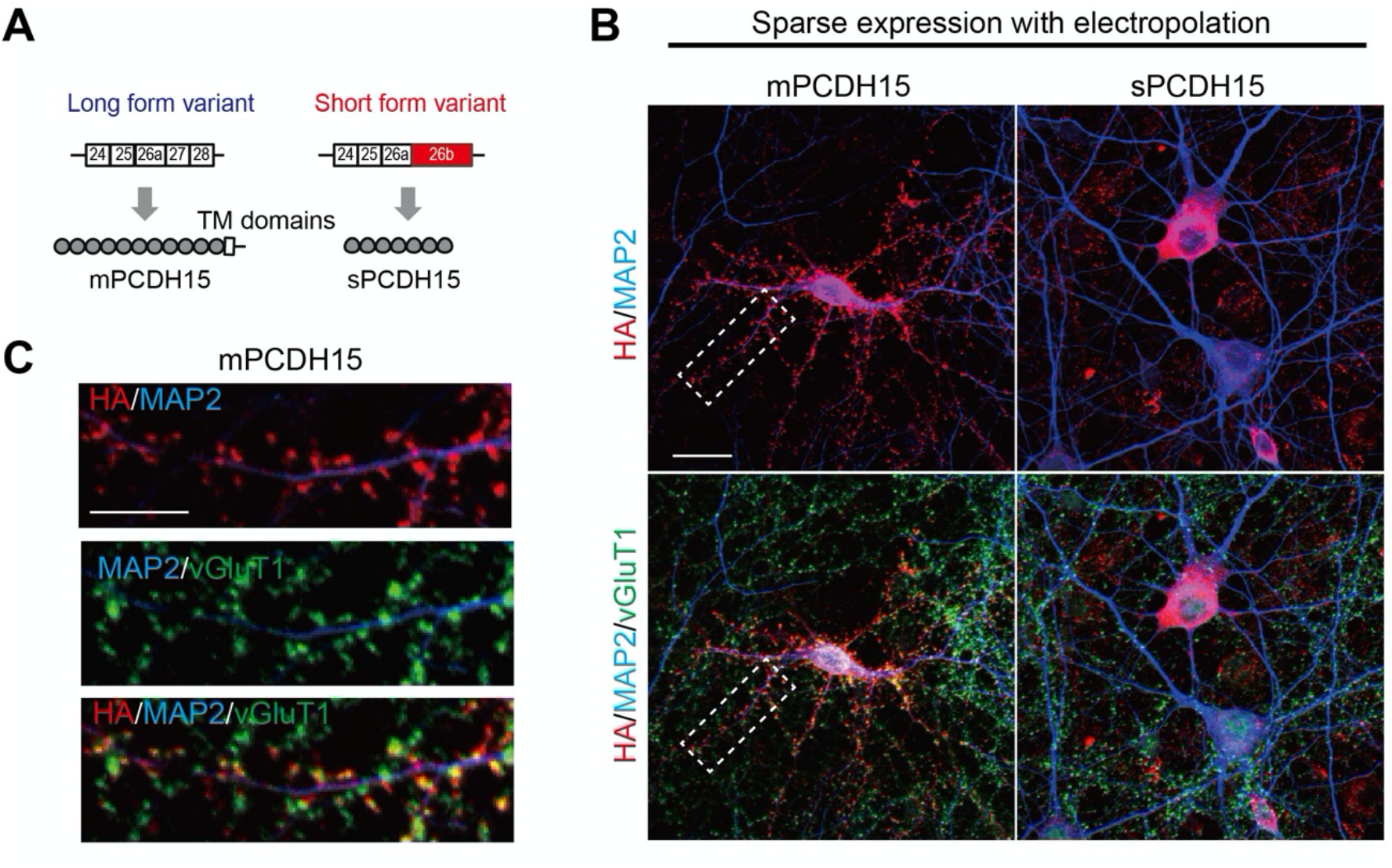
Aberrant ALE splicing disrupts PCDH15 localization into excitatory postsynapses. (A) Schematic illustration of the ALE selection of *Pcdh15*. The insertion of exon 26b causes the truncation of the large coding sequence at the C’ terminal and transmembrane (TM) domain and expresses the soluble form (sPCDH15) instead of the membrane-bound type (mPCDH15). (B) Immunostaining of the subcellular localization of mPCDH15 (left) and sPCDH15 (right) in cultured cortical neurons (at DIV14) after electroporation with HA-tagged mPCDH15 or sPCDH15 lentiviral vectors at DIV4. VGluT1 and MAP2 stained excitatory synaptic boutons and the overall morphology of neurons, respectively. (C) High magnification images of the area surrounded by dashed rectangles in (B). Scale bars = 10 μm in (B) and (C).

We next attempted to assess the subcellular localization of PCDH15 in cultured neurons with several antibodies against PCDH15, but immunoreactivity was not reliably detected with these antibodies. Therefore, we sparsely expressed mPCDH15 and sPCDH15 tagged with hemagglutinin (HA) at the C’ terminal in cultured cortical neurons and evaluated subcellular localization with an anti-HA antibody. The immunoreactivity of mPCDH15-HA was robustly accumulated on the spine heads of the neurons (Figure 4B). The localization at the spines was confirmed by co-staining with a vesicular glutamate transporter 1 (VGluT1) antibody, which is an excitatory synaptic marker (Figure 4C), suggesting that mPCDH15 is abundantly localized into excitatory post-synaptic neurons. Most of the sPCDH15-HA was retained in the soma and not inserted into the postsynapses (Figure 4B). Thus, the ALE selection of *Pcdh15* dramatically affected its synapse localization in cortical neurons.

### Aberrant ALE selection of Pcdh15 influences synapse development

Given that ALE alterations in *Pcdh15* alter the synaptic localization of PCDH15, we examined the effects of ALE alterations in *Pcdh15* on synapse development and compared them with *Pcdh15* mutants and controls (mock). The CRISPR/Cas9 system was used to generate *Pcdh15* mutations by removing the constitutive exon 6, which yielded a frameshift truncation mutation (Figure S3A). The *Pcdh15* mutation (Δex6) mimicked a rare variant found in Usher syndrome patients [50-bp del NM_001384140.1(PCDH15): c.475-2204_594+1766 del] (19). qPCR analyses revealed that exon 6 was removed in *Pcdh15*Δex6 (*gPcdh15*) neurons and increased with the expression of *sPcdh15* in cortical neuron cultures (Figure S3B). We first assessed the effects of the Δex6 mutation and ALE alterations in *Pcdh15* on excitatory synapses by analyzing the numbers and sizes of excitatory synapses in the Δex6 mutant and sPCDH15-expressing neuron cultures. Immunostaining with anti-PSD95 and anti-VGluT1 antibodies, which are excitatory post-synaptic markers, showed that the number of synapses was significantly reduced and spine size was significantly increased in the Δex6 mutant cultures (Figures 5A and 5B, S3C). The expression of sPCDH15 significantly reduced the number of synapses mimicking the Δex6 mutant phenotype; however, the spine size did not significantly change (Figure 5A and 5C). In addition, there were no significant differences in the densities of the neurons (MAP2^+^/DAPI^+^ cells) (Figure S3D) and transcript levels of the synaptic markers (Figure S3E) between the groups, indicating that the altered number of excitatory synapses was not caused by the reduced number of neurons or transcriptional changes.

**Figure 5:**
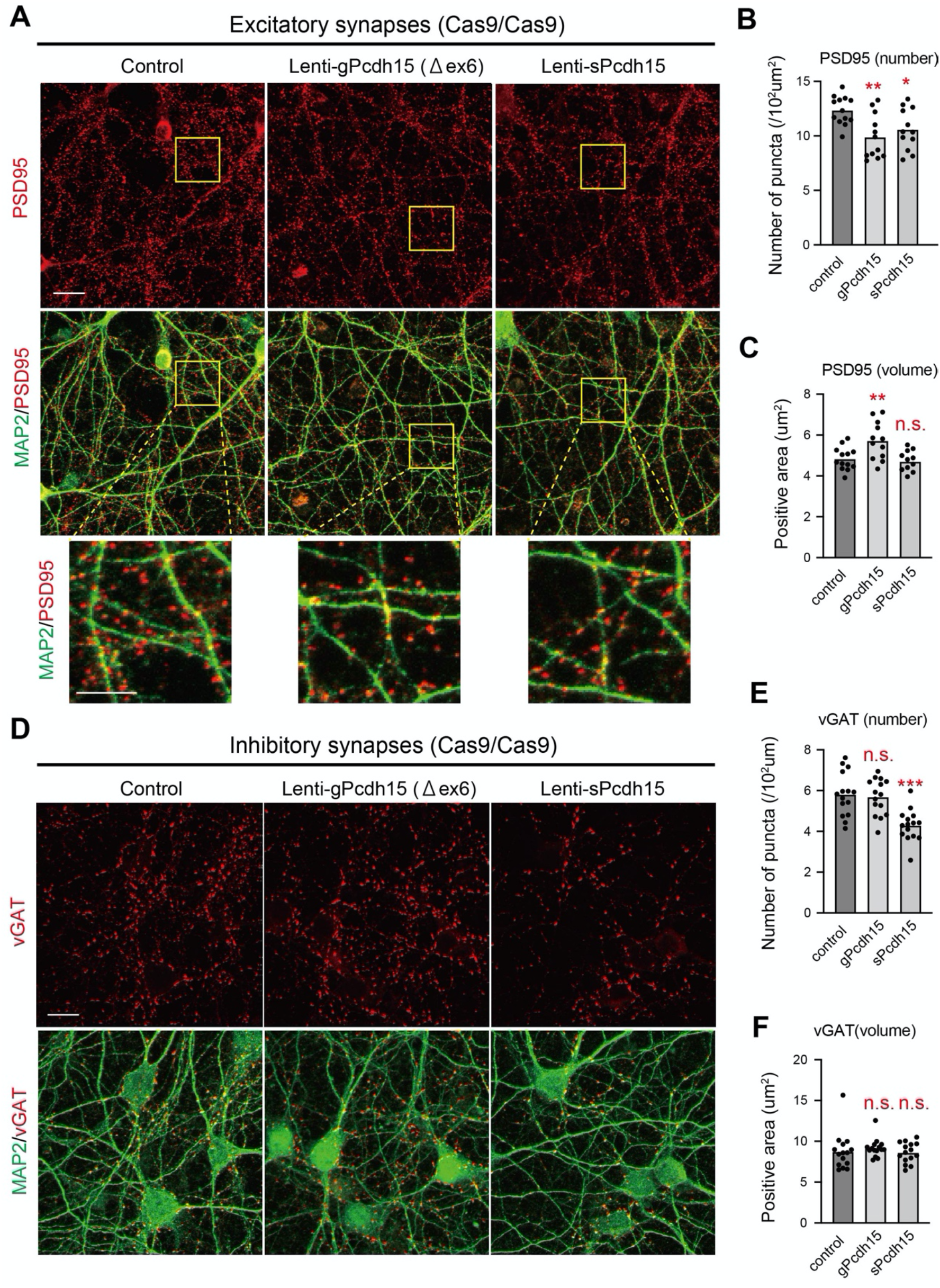
The aberrant ALE isoform expression of *Pcdh15* influences excitatory and inhibitory synapses while the *Pcdh15* Δex6 mutation preferentially affects excitatory synapses. (A) Immunostaining of Cas9-expressing cortical neuronal cultures transduced with mock lentiviral vectors (control, left), lentiviral vectors carrying gRNAs to delete Pcdh15 (Δex6 mutation) (*gPcdh15*, middle), and *sPcdh15*-expressing vectors (sPcdh15, right) with the neuronal markers PSD95 and MAP2. The upper panels show PSD95-expressing puncta. The middle panels show an overlay between PSD95 and MAP2. The lower panel shows high magnification images of the area surrounded by dashed rectangles in the middle panels. (B, C) Density (B) and volume (C) of excitatory synapses treated with lentiviral vectors carrying the control, *gPcdh15,* and *sPcdh15*. n = 13, 13, and 12 fields for the control, *gPcdh15*, and *sPcdh15*, respectively. A one-way ANOVA was followed by Tukey’s multiple comparisons test. (B) Synaptic number. One-way ANOVA [F (2, 34) = 6.609, P = 0.004] (C) Synaptic volume. One-way ANOVA [F (2, 34) = 7.979, P = 0.0014]. (D) Immunostaining of Cas9-expressing cortical neuronal cultures transduced with mock lentiviral vectors (control, left), lentiviral vectors carrying gRNAs to delete *Pcdh15* (Δex6 mutation) (*gPcdh15*, middle), and *sPcdh15*-expressing vectors (*sPcdh15*, right) with the neuronal markers VGAT and MAP2. The upper panels show VGAT-expressing puncta. The middle panels show an overlay between VGAT and MAP2. (E, F) Density (E) and volume (F) of inhibitory synapses treated with lentiviral vectors carrying the control, g*Pcdh15,* and *sPcdh15*. n = 15 fields per group. (E) synaptic number. One-way ANOVA [F (2, 42) = 12.75, P < 0.0001] (F) synaptic volume. Scale bars = 10 μm in (A) and (D).

We next analyzed the influences of the Δex6 mutation and ALE alterations in *Pcdh15* on inhibitory synapses. Immunostaining with an anti-vesicular GABA transporter (VGAT) antibody, which is an inhibitory post-synaptic marker, showed that the expression of sPCDH15 significantly reduced the number of inhibitory synapses, whereas they did not significantly alter in the Δex6 mutant. The spine size did not significantly change in either group. Therefore, while the Δex6 mutation of *Pcdh15* yielded slight effects on the excitatory synapses, the expression of sPCDH15 likely influenced both the excitatory and inhibitory synapses.

### Preferential effects of aberrant ALE isoforms of Pcdh15 on inhibitory synapses

To confirm the effects of sPCDH15 on neuronal synapses, we measured excitatory and inhibitory neurotransmitters released in the expression cultures. Liquid chromatography–mass spectrometry (LC–MS) was used to quantify the spontaneous and evoked releases of neurotransmitters before and after pharmacological stimulations with 4-AP, respectively. While the spontaneous and evoked releases of glutamate were slightly reduced in the sPCDH15- expressing cultures (Figure 6A), those of GABA were dramatically reduced (Figure 6B). The releases of both neurotransmitters were significantly enhanced in the evoked cultures compared to the unstimulated cultures (Figure 6A and 6B), which confirmed the functional synaptic response to depolarized stimulation. These data suggest that sPCDH15 has a more potent effect on inhibitory synapses than excitatory ones.

**Figure 6:**
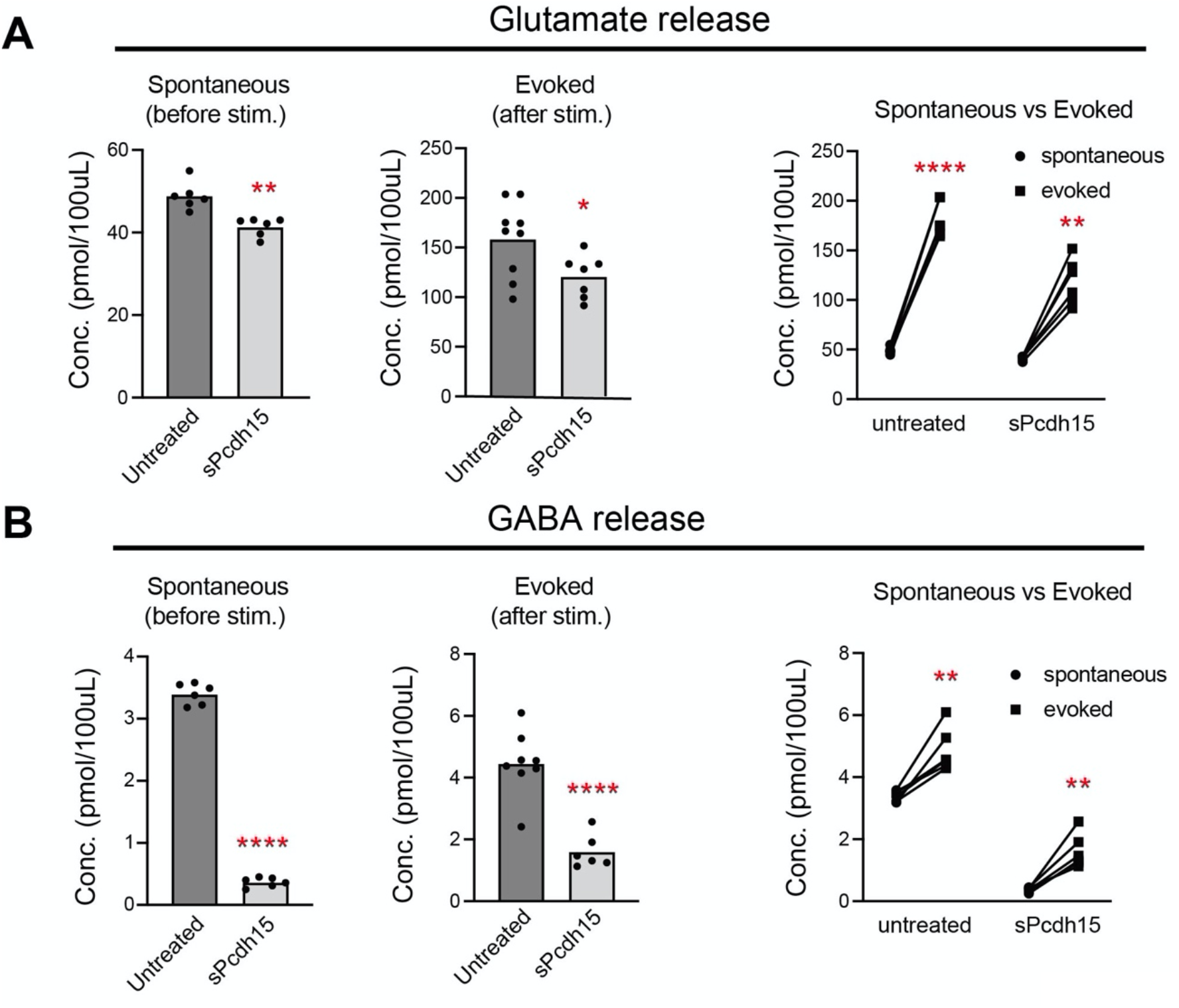
The expression of sPCDH15 preferentially impairs inhibitory synapses. (A, B) Concentrations of released glutamate (A) and GABA (B) in cortical neurons transduced with lentivirus-expressing mock genes (control) or *sPcdh15* (*sPcdh15)*. The neurotransmitter level was measured upon a steady-stead level (spontaneous, left panel) and after the neurons had been depolarized with 4-AP (evoked, middle panel). The right panels show paired comparisons before and after neuronal stimulation in both groups. n = 8 and 6 cultures (control and sPcdh15), unpaired *t*-test. The functional response to stimulation was checked using a paired comparison before and after stimulation (right panels). n = 6 cultures per group, paired *t*-test.

We further explored the preferential effect of sPCDH15 on inhibitory synapses. sPCDH15 was not localized at synaptic boutons but resided within the soma upon transient expression with *in vitro* electroporation (Figure 4A and 4B). However, when sPCDH15 was expressed for a long-term period with lentiviral infection (DIV7–14 for 7 days), immunoreactivity was partially colocalized with the anti-VGAT antibody in addition to the expression in the soma (Figure 7A and 7B). This implies that the aberrant ALE selection of sPCDH15 causes the mislocalization of PCDH15 into inhibitory synapses. To understand the mechanisms underlying the preferential action of sPCDH15 on inhibitory synapses, we examined the effects of sPCDH15 on the synaptogenic activity of inhibitory neurons *in vitro*. To achieve this, we analyzed the effects of sPCDH15 on the assembly of inhibitory synapses induced by neuroligin- 2 (NL2), which is an inhibitory synapse organizer in the nervous system, in a neuron-fibroblast co-culture assay (Figure 7C). NL2 can strongly induce artificial presynaptic contacts for both excitatory and inhibitory neurons but preferentially induces inhibitory synapses more than other synapse organizers (26). The expression of NL2 in human embryonic kidney (HEK293T) cells significantly formed the assembly of both VGluT1^+^ and VGAT^+^ presynapses. In addition, the expression of mPCDH15 did not induce any synapse formation (Figure 7D). However, when sPcdh15 was co-expressed with NL2, the assembly of VGAT^+^, but not VGluT1^+^, synapses were significantly attenuated (Figure 7D and 7E), indicating that sPCDH15 selectively prevents the formation of inhibitory synapses *in vitro*.

**Figure 7:**
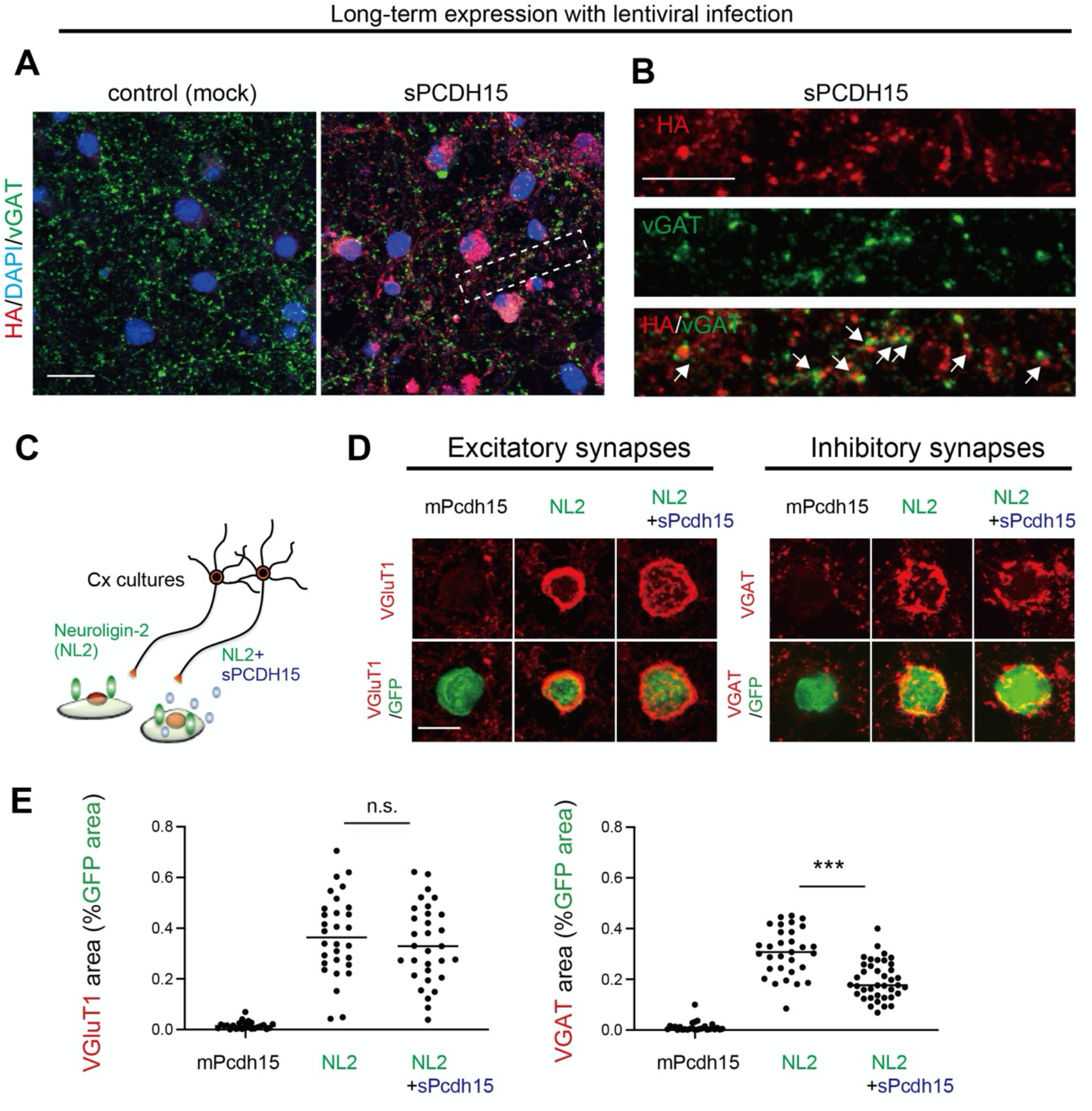
Selective effects of the aberrant ALE isoform of PCDH15 on the formation of NL2-induced inhibitory synapses. (A) Immunostaining of the subcellular localization of sPCDH15 in cultured cortical neurons transduced with lentiviral vectors expressing HA-tagged sPCDH15. VGAT and DAPI stained inhibitory synaptic boutons and neuronal nuclei, respectively. sPcdh15 was partially detected in the boutons of VGAT-positive inhibitory synapses. (B) High magnification images of the area surrounded by dashed rectangles in (A). The upper, middle, and lower panels show the staining of sPCDH15 alone, VGAT alone, and overlay, respectively. (C) Cartoon showing the artificial synapse formation assay. NL2-HA was expressed into HEK293T cells with and without sPcdh15-HA and co-cultured with cortical neurons. (D) Immunostaining of HEK293T cells expressing Pcdh15, NL2, and NL2+Pcdh15 with the presynaptic markers VGluT1 (excitatory synapses, left panel) and VGAT (inhibitory synapses, right panel). The overall morphology of co-cultured HEK293T cells was visualized with a GFP. (E) The quantification of synapse assembly described in (D) was achieved by measuring the VGLUT1 (left panel) or VGAT areas (right panel) relative to the GFP area. VGluT1: [n = 25, 30, and 31 cells for mPcdh15, NL2, and NL2+sPcdh15, respectively]. One-way ANOVA [F (2, 83) = 57.19, P < 0.0001]. VGAT: [n = 27, 31, and 39 cells for mPcdh15, NL2, and NL2+sPcdh15, respectively]. One-way ANOVA [F (2, 94) = 122.8, P < 0.0001], followed by Tukey’s multiple comparisons test. Scale bars = 10 μm in (A) and (B), 5 μm in (D).

## Discussion

### The control of SAM68-mediated ALE selection by distinct mechanisms in the nervous system

The ALE isoforms of many genes are spatially and temporally altered in mammalian tissues. Recently, it was found that SAM68 is expressed in a tissue-specific manner and is required for the spatial control of ALE isoforms (10–12). Our study further demonstrated that the ALE splicing of several transcripts is regulated by SAM68 in a dose-dependent manner and can be dynamically exerted by external regulations. SAM68 is further modulated by several cellular signals that alter the AS of various targets (23,24). Given that SAM68 regulates neuronal activity-dependent AS through the CaMK pathway (22), we speculated whether SAM68- regulated ALE selection is affected similarly. As expected, various forms of SAM68-regulated ALE selection were modulated upon spontaneous activity and further controlled by cellular signals. Additionally, the ALE of *Pcdh15*, but not *Il1rap*, was partially affected by the depletion of spontaneous activity and CaMK signals (Figure 2), which may be mechanisms of spatiotemporal ALE splicing in the nervous system.

In contrast to *Pcdh15*, the ALE of *Il1rap* was likely dependent on U1 snRNP (Figure 3). This is conceivable because U1 motif-like sequences were located around the cryptic PASs of *Il1rap* and absent around those of *Pcdh15*. The regulation of SAM68 targets by U1 snRNP was not surprising because recent studies have suggested that SAM68 synergistically regulates the 3’-end processing of target transcripts with U1 snRNP complexes (11,13) by protecting pre-mRNAs from drastic premature termination by cleavage and PASs in introns (14,25). Thus, SAM68-regulated ALE splicing appears to be complex and likely depends on targets.

Given these findings, it is conceivable that the cooperative action of U1 snRNP and SAM68 allows for the regulation of the ALE to be more stable and less susceptible to external regulation, whereas the regulation of the ALE solely by SAM68 is likely to make it more prone to external effects. Thus, we suggest that the differences in the regulation of the ALE by SAM68 between target transcripts are likely determined by the presence or absence of co-factors such as U1 snRNP. However, the detailed mechanisms underlying the dependency of target transcripts on neuronal activity and cellular pathways remain to be addressed in future studies.

### Maintenance and synaptic role of proper ALE splicing in Pcdh15 by SAM68

Mutations, CNVs, and SNPs of *Pcdh15* are implicated in autism, bipolar disorder, schizophrenia, and Usher syndrome (17,18) (27) (20) (21), but its expression, localization, and functions in the central nervous system remain unclear. We found that PCDH15 is highly localized in excitatory postsynapses and that the pathogenic loss-of-function mutation caused a reduction in the excitatory synapses (Figures 4 and 5). Given that a recent study further showed that the human induced pluripotent stem cell-derived neurons from two patients with bipolar disorder and a small deletion in the *Pcdh15* gene exhibited decreasing synapse numbers (28), PCDH15 may play a pivotal role in various aspects of synaptic functions. Dysregulated *Pcdh15* expression may therefore be implicated in the pathologies of several psychiatric disorders. Similarly, we found that the expression of the aberrant ALE isoform, sPCDH15, slightly disrupted the formation and function of excitatory synapses, thereby mimicking the pathogenic loss-of-function mutation. Given that sPCDH15 is secreted into cultured media (10), the impairment may be due to the competitive effect at excitatory synapses.

The expression of sPCDH15 was further found to preferentially influence inhibitory synapses more than excitatory ones (Figures 5, 6). Since PCDH15 can exist at excitatory postsynapses, the effects of sPCDH15 on inhibitory synapses may be surprising. However, we observed that sPCDH15 was widely dispersed following a lengthy expression period with lentiviral infection. Immunoreactivity was observed around VGAT-positive puncta, which may explain the effects of sPCDH15 on inhibitory neurons (Figure 7A). In addition, sPCDH15 significantly prevented NL2-induced artificial synapse formation (Figure 7B and 7C). Mislocalized sPCDH15 may therefore negatively affect inhibitory synapses. This finding is somewhat consistent with recent studies in which a clustered PCDH, γ-protocadherin, interacted with NL-1/2 (29) (30) and negatively regulated inhibitory synapse density *in vivo* (30).

Given that altered 3’UTR exons in *Sam68* ^KO^ brains include a significant number of transcripts encoding transmembrane or secreted proteins other than PCDH15 and IL1RAP with neuronal functions, it is conceivable that ALE selection by SAM68 may be involved in a plethora of neuronal and synaptic functions.

## Experimental procedures

### Antibodies and plasmids

The following commercially available antibodies were used for the immunostaining analyses: mouse anti-HA (clone HA-7, Sigma-Aldrich, St. Louis, MO, USA), mouse anti-PSD95 (clone K28/43, NeuroMab, Davis, CA, USA), rabbit anti-MAP2 (Sigma-Aldrich), rabbit anti-VGluT1 (#1353303, Synaptic Systems, Göttingen, Germany), and guinea pig anti-VGAT (#676780, Calbiochem, San Diego, CA, USA). Secondary antibodies with minimal interspecies cross-reactivity conjugated to Alexa Fluor dyes (Molecular Probes, Eugene, OR, USA) were used for visualization in the immunostaining analyses. To visualize the overall morphology of transfected HEK293T cells in artificial synapse formation assay (see below), we used previously-described expression vectors comprising green fluorescent protein (GFP), NL2 (containing the A site), mPcdh15, and sPcdh15 (pMAX-GFP, pCAGGS-NL2A-HA, pCAGGS- mPcdh15-HA, and pCAGGS-sPcdh15-HA) (31) (10).

### Neuronal cell culture and artificial synapse formation assays

Cortical neuron cultures containing ganglionic eminence were prepared as previously described (32). The cultures were prepared from pops of ICR or Cas9-expressing mouse pups on embryonic day 15 (E15). The tissues were dissociated with 0.05% trypsin (Sigma-Aldrich) in the presence of DNase I (Roche Applied Science, Penzberg, Germany) for 10 min at 37 °C. After the cells had been dissociated, trypsin was inactivated with a soybean trypsin inhibitor (Sigma-Aldrich). The cells were then plated into poly-D-ornithine-coated dishes (2.0 × 10^5^/cm^2^) and maintained for 15 days in Neurobasal Medium (Invitrogen, Waltham, MA, USA) containing 2% B27 supplement, 2 mM of Glutamax, and penicillin/streptomycin (Invitrogen). For the pharmacological experiments, TTX, Bic, STO609, KN93, U0126, and PP2 purchased from TOCRIS (Bristol, UK) or Sigma-Aldrich were added 1 day before harvesting. All procedures related to the care and treatment of animals were carried out in strict accordance with the Guide for the Care and Use of Laboratory Animals of Tokai University. All mice were maintained under specific pathogen-free conditions at the Laboratory Animal Center, Tokai University. The experimental protocol was approved by the Institutional Animal Care and Use Committee of Tokai University (permit number 224036). All surgeries were performed under sodium pentobarbital anesthesia with efforts made to minimize animal suffering.

For the immunostaining analyses, cultured neurons were fixed with 4% paraformaldehyde in ice-cold phosphate-buffered saline (PBS) for 15 min. The neurons were then permeabilized with PBS containing 0.15% TritonX-100 for 15 min at room temperature and incubated with blocking solution (5% normal goat serum in PBS) for at least 30 min at room temperature. They were then incubated with the primary antibodies for 24 h at 4 °C. The appropriate secondary antibodies conjugated to Alexa 546 or 488 (goat; 1:1000) (Life Technology, Carlsbad, CA, USA) were used for visualization.

For the artificial synapse formation assay, HEK293T cells co-expressing trans-synaptic NRX receptors with GFP were plated on the cortical neuron cultures at DIV14 (1–2 x 10^4^ cells/cm^2^) and immunostained with the presynaptic markers and anti-VGluT1 and anti-VGAT antibodies.

### Image acquisition and analysis

Confocal images of the neuronal cell cultures were captured using an LSM700 confocal system (Zeiss, Göttingen, Germany). The original images were analyzed using ImageJ (NIH, Bethesda, MD, USA). The number and size (area) of the synapse marker-positive puncta defined with proper thresholding (top 2.5–5.0% of the display range) were quantified. The morphologies of the neurons were visualized by staining with the neuronal marker MAP2. Nuclei were visualized with DAPI stains.

Quantification in the artificial synapse formation assay was performed as previously described [22, 24]. In brief, after the appropriate threshold had been set, the immunoreactive areas on surface of the HEK293T cells were measured. Approximately 20–40 cells from more than 10 separate fields per culture were quantified per group. The morphologies of the HEK293T cells were visualized by co-expressing GFP.

### RNA isolation, semi-qPCR, and RT-qPCR assays

For the qPCR analyses, RNA was harvested from cultured neurons using Nucleospin RNA XS (TaKaRa, Tokyo, Japan) or RNAiso Plus reagent (TaKaRa), followed by the removal of contaminating DNA using Turbo DNA-free (RNase-free DNase; Ambion, Austin, TX, USA). Total RNA (1–2 µg) was reverse transcribed using random hexamers and the PrimeScript™ 1st strand cDNA Synthesis Kit (TaKaRa). RT-qPCR was performed using a StepOnePlus qPCR system (Applied Biosystems, Waltham, MA, USA) with Power SYBR Green PCR Master Mix (Applied Biosystems) and the comparative CT method. For relative quantification using qRT-PCR, transcript levels were normalized to that of *Gapdh*. The oligonucleotide primers used for the qPCR are listed in Table S1.

For the abundance ratio of the short form (SF) to long form (LF), the percentage of the SF variant was largely estimated from the CT value (CT^SF^) and directly compared to that of the LF (CT^LF^) at the same threshold set for the CT value. The RQ^LF^+RQ^SF^ values for the total transcript levels were set to 100%.

### Lentiviral production and generation of vectors for *in vitro* genome editing

The procedures for the lentiviral production have been previously described (Suzuki et al., 2017). pCL20c vectors were designed under the control of the murine stem cell virus (MSCV) promoter (33). The viral vector was produced by co-transfecting HEK293T cells with a mixture of four plasmids using a calcium phosphate precipitation method. The four-plasmid mixture consisted of 6 µg of pCAG-kGP1R, 2 µg of pCAG-4RTR2, 2 µg of pCAG-VSV-G, and 10 µg of the vector plasmid pCL20c (pCL20c-MSCV-sPcdh15-HA-IRES-EGFP). The medium containing vector particles was harvested 40 h after transfection. Media samples were concentrated by centrifugation at 15,000 x g for 5–7 h. Viral samples were then suspended in Neurobasal Medium, frozen in aliquots, and stored at –80 °C until further use. After assessing the titer in the HEK293T cells, the appropriate amount of lentivirus was transduced into cultured neurons 5–7 days before harvesting the cells.

For the design of the lentiviral vectors for the *in vitro* genome editing, the sequences of gRNAs were selected based on CHOPCHOP (https://chopchop.cbu.uib.no) (>Rank5) (see Table S1). The Lentiviral CRISPR/Cas9 System (System Biosciences LLC, Palo Alto, CA, USA) was then used according to the manufacturer’s protocol to construct the vector, and the oligonucleotides of the gRNAs were inserted into the U6 promoter-driven lentiviral vector (CASLV511PA-G).

### AMOs and electroporation

The sequences of U1 and the control AMOs were used as previously described (14). The AMOs were purchased from Gene Tools, LLC (Philomath, OR, USA). AMO transfection was performed using the Neon Transfection System (Life Technologies, Carlsbad, CA, USA). The cells (2×10^7^ cells) were resuspended in 100 µL of the resuspension buffer containing the AMOs. They were then electroporated and plated onto 12- or 24-well culture dishes.

Electroporation was performed using the following parameters based on manufactural protocol: 1500 V (pulse voltage), 10 msec (pulse width), and three pulses.

### Measurement of neurotransmitter release by LC–MS

The culture supernatant was collected, dried using a SpeedVac vacuum concentrator (Thermo Fischer Scientific, Waltham, MA, USA), and reconstituted in 100 µL of methanol, of which 10 µL were injected into an LC-MS. A quantitative analysis of the neurotransmitters was performed using an LCMS-8050 triple quadrupole mass spectrometer coupled to the UHPLC-Nexera system (Shimadzu, Kyoto, Japan). The chromatographic condition of the neurotransmitters was optimized using modifications described in previously published procedures (34,35). The samples were separated in a hydrophilic interaction liquid chromatography mode using a 2.1 × 150 mm, 3.5 µm iHILIC Fusion(P) zwitterionic column (HILICON AB, Umeå, Sweden). Mobile phase A consisted of 100 mM of ammonium bicarbonate containing 5 µM of medronic acid (pH 9.0). Mobile phase B included 90% LC–MS-grade acetonitrile containing 10 mM of ammonium bicarbonate and 5 µM of medronic acid (pH 9.0). Linear gradient elution was achieved using the following procedure: 0–2 min, 95% B; 2–10 min, 95-5% B; 10–12.5 min, 5% B. The gradient was then restored to the initial composition (95%) for equilibration before the next run. The total flow rate was 2.0 mL/min. Selected reaction monitoring in positive electrospray ionization mode was performed to detect GABA (m/z 104.15 > 87.10 for quantitation, m/z 104.15 > 69.10 for identification) and glutamic acid (m/z 148.15 > 84.10 for quantitation, m/z 148.15 > 56.10 for identification).

### Statistical analysis

GraphPad Prism 5 (GraphPad Software, Inc., La Jolla, CA, USA) was used for the statistical analyses. D’Agostino-Pearson or Shapiro-Wilk normality tests were used to ensure the data had been normally distributed. Comparisons between two groups were made using Student’s *t*-tests, and those between three or more groups were made using an analysis of variance (ANOVA) followed by Tukey’s or Dunnett’s tests for pairwise comparisons, unless otherwise stated. Data were presented as individual data points and means. Significance levels were indicated as follows: ****P < 0.0001; ***P < 0.001; **P < 0.01; *P < 0.05.

## Data availability

The data that support the findings of this study are available from the corresponding author (T.I.) upon reasonable request. The RNA-seq data can be accessible at the DDBJ database (6781), with accession number of DRA6781, with bioproject accession number of PRJDB6781.

## Supporting information

This article contains supporting information.

## Acknowledgments

We thank all the members of the Support Center for Medical Research and Education at Tokai University for their experimental support and maintenance of the experimental animals. We appreciate Prof. Ullich Müller (Scripps Res. Inst., USA) for kindly providing materials for Pcdh15.

## Author contributions

T.I. conceived and designed the experiments. T.I., M.D., M.I., A.T., N.A., S.S., M.T., and Y.I. performed the experiments and analyzed the data; T.I. and M.D. wrote the paper.

## Funding and additional information

This study was supported by research grants from the JSPS KAKENHI (grant no. 15H04277 and 20H03344) (to T.I.), the Takeda Science Foundation (to T.I.), Naito Foundation, Novartis Foundation (to T.I.), and Kanehara Memorial Foundation (to T.I.).

## Abbreviations

3’UTR: 3’ untranslated region
ALE: alternative last exon
Pcdh15: protocadherin-15
sPCDH15: soluble form of PCDH15
APA: alternative polyadenylation
U1 snRNP: U1 small nuclear ribonucleoprotein
CNV: copy number variation
SNP: single nucleotide polymorphism
RNA-seq: RNA sequencing
Il1rap: interleukin 1-receptor accessory protein
Cp: ceruloplasmin
Lrrcc1: leucine-rich repeated coiled-coil protein 1
AS: alternative splicing
Bic: bicuculine
TTX: tetrodotoxin
PAS: polyadenylation site
AMO: antisense morpholino oligonucleotide
qPCR: quantitative PCR
mPCDH15: membrane-bound form of PCDH15
VGluT1: vesicular glutamate transporter 1
VGAT: vesicular GABA transporter
LC–MS: liquid chromatography–mass spectrometry
NL2: neuroligin-2
PBS: phosphate-buffered saline
MSCV: murine stem cell virus
ANOVA: analysis of variance

## References

1. Tian, B., Hu, J., Zhang, H., and Lutz, C. S. (2005) A large-scale analysis of mRNA polyadenylation of human and mouse genes. Nucleic Acids Res 33, 201–212

2. Miura, P., Shenker, S., Andreu-Agullo, C., Westholm, J. O., and Lai, E. C. (2013) Widespread and extensive lengthening of 3’ UTRs in the mammalian brain. Genome Res 23, 812–825

3. Reyes, A., and Huber, W. (2018) Alternative start and termination sites of transcription drive most transcript isoform differences across human tissues. Nucleic Acids Res 46, 582–592

4. Lianoglou, S., Garg, V., Yang, J. L., Leslie, C. S., and Mayr, C. (2013) Ubiquitously transcribed genes use alternative polyadenylation to achieve tissue-specific expression. Genes Dev 27, 2380–2396

5. Sandberg, R., Neilson, J. R., Sarma, A., Sharp, P. A., and Burge, C. B. (2008) Proliferating cells express mRNAs with shortened 3’ untranslated regions and fewer microRNA target sites. Science 320, 1643–1647

6. Taliaferro, J. M., Vidaki, M., Oliveira, R., Olson, S., Zhan, L., Saxena, T., Wang, E. T., Graveley, B. R., Gertler, F. B., Swanson, M. S., and Burge, C. B. (2016) Distal Alternative Last Exons Localize mRNAs to Neural Projections. Mol Cell 61, 821–833

7. Gruber, A. J., and Zavolan, M. (2019) Alternative cleavage and polyadenylation in health and disease. Nat Rev Genet 20, 599–614

8. Tian, B., and Manley, J. L. (2017) Alternative polyadenylation of mRNA precursors. Nat Rev Mol Cell Biol 18, 18–30

9. Holt, C. E., and Schuman, E. M. (2013) The central dogma decentralized: new perspectives on RNA function and local translation in neurons. Neuron 80, 648–657

10. Iijima, Y., Tanaka, M., Suzuki, S., Hauser, D., Tanaka, M., Okada, C., Ito, M., Ayukawa, N., Sato, Y., Ohtsuka, M., Scheiffele, P., and Iijima, T. (2019) SAM68-specific splicing is required for proper selection of alternative 3’ UTR isoforms in the nervous system. iScience 22, 318–335

11. Naro, C., Pellegrini, L., Jolly, A., Farini, D., Cesari, E., Bielli, P., de la Grange, P., and Sette, C. (2019) Functional Interaction between U1snRNP and Sam68 Insures Proper 3’ End Pre-mRNA Processing during Germ Cell Differentiation. Cell Rep 26, 2929–2941 e2925

12. La Rosa, P., Bielli, P., Compagnucci, C., Cesari, E., Volpe, E., Farioli Vecchioli, S., and Sette, C. (2016) Sam68 promotes self-renewal and glycolytic metabolism in mouse neural progenitor cells by modulating Aldh1a3 pre-mRNA 3’-end processing. Elife 5

13. Subramania, S., Gagne, L. M., Campagne, S., Fort, V., O’Sullivan, J., Mocaer, K., Feldmuller, M., Masson, J. Y., Allain, F. H. T., Hussein, S. M., and Huot, M. E. (2019) SAM68 interaction with U1A modulates U1 snRNP recruitment and regulates mTor pre-mRNA splicing. Nucleic Acids Res 47, 4181–4197

14. Kaida, D., Berg, M. G., Younis, I., Kasim, M., Singh, L. N., Wan, L., and Dreyfuss, G. (2010) U1 snRNP protects pre-mRNAs from premature cleavage and polyadenylation. Nature 468, 664–668

15. Kim, S. Y., Yasuda, S., Tanaka, H., Yamagata, K., and Kim, H. (2011) Non-clustered protocadherin. Cell Adh Migr 5, 97–105

16. Ahmed, Z. M., Riazuddin, S., Bernstein, S. L., Ahmed, Z., Khan, S., Griffith, A. J., Morell, R. J., Friedman, T. B., Riazuddin, S., and Wilcox, E. R. (2001) Mutations of the protocadherin gene PCDH15 cause Usher syndrome type 1F. Am J Hum Genet 69, 25–34

17. Alagramam, K. N., Yuan, H., Kuehn, M. H., Murcia, C. L., Wayne, S., Srisailpathy, C. R., Lowry, R. B., Knaus, R., Van Laer, L., Bernier, F. P., Schwartz, S., Lee, C., Morton, C. C., Mullins, R. F., Ramesh, A., Van Camp, G., Hageman, G. S., Woychik, R. P., and Smith, R. J. (2001) Mutations in the novel protocadherin PCDH15 cause Usher syndrome type 1F. Hum Mol Genet 10, 1709–1718

18. Georgieva, L., Rees, E., Moran, J. L., Chambert, K. D., Milanova, V., Craddock, N., Purcell, S., Sklar, P., McCarroll, S., Holmans, P., O’Donovan, M. C., Owen, M. J., and Kirov, G. (2014) De novo CNVs in bipolar affective disorder and schizophrenia. Hum Mol Genet 23, 6677–6683

19. Carss, K. J., Arno, G., Erwood, M., Stephens, J., Sanchis-Juan, A., Hull, S., Megy, K., Grozeva, D., Dewhurst, E., Malka, S., Plagnol, V., Penkett, C., Stirrups, K., Rizzo, R., Wright, G., Josifova, D., Bitner-Glindzicz, M., Scott, R. H., Clement, E., Allen, L., Armstrong, R., Brady, A. F., Carmichael, J., Chitre, M., Henderson, R. H. H., Hurst, J., MacLaren, R. E., Murphy, E., Paterson, J., Rosser, E., Thompson, D. A., Wakeling, E., Ouwehand, W. H., Michaelides, M., Moore, A. T., Consortium, N. I.-B. R. D., Webster, A. R., and Raymond, F. L. (2017) Comprehensive Rare Variant Analysis via Whole-Genome Sequencing to Determine the Molecular Pathology of Inherited Retinal Disease. Am J Hum Genet 100, 75–90

20. Kushima, I., Aleksic, B., Nakatochi, M., Shimamura, T., Shiino, T., Yoshimi, A., Kimura, H., Takasaki, Y., Wang, C., Xing, J., Ishizuka, K., Oya-Ito, T., Nakamura, Y., Arioka, Y., Maeda, T., Yamamoto, M., Yoshida, M., Noma, H., Hamada, S., Morikawa, M., Uno, Y., Okada, T., Iidaka, T., Iritani, S., Yamamoto, T., Miyashita, M., Kobori, A., Arai, M., Itokawa, M., Cheng, M. C., Chuang, Y. A., Chen, C. H., Suzuki, M., Takahashi, T., Hashimoto, R., Yamamori, H., Yasuda, Y., Watanabe, Y., Nunokawa, A., Someya, T., Ikeda, M., Toyota, T., Yoshikawa, T., Numata, S., Ohmori, T., Kunimoto, S., Mori, D., Iwata, N., and Ozaki, N. (2017) High-resolution copy number variation analysis of schizophrenia in Japan. Mol Psychiatry 22, 430–440

21. Kushima, I., Nakatochi, M., Aleksic, B., Okada, T., Kimura, H., Kato, H., Morikawa, M., Inada, T., Ishizuka, K., Torii, Y., Nakamura, Y., Tanaka, S., Imaeda, M., Takahashi, N., Yamamoto, M., Iwamoto, K., Nawa, Y., Ogawa, N., Iritani, S., Hayashi, Y., Lo, T., Otgonbayar, G., Furuta, S., Iwata, N., Ikeda, M., Saito, T., Ninomiya, K., Okochi, T., Hashimoto, R., Yamamori, H., Yasuda, Y., Fujimoto, M., Miura, K., Itokawa, M., Arai, M., Miyashita, M., Toriumi, K., Ohi, K., Shioiri, T., Kitaichi, K., Someya, T., Watanabe, Y., Egawa, J., Takahashi, T., Suzuki, M., Sasaki, T., Tochigi, M., Nishimura, F., Yamasue, H., Kuwabara, H., Wakuda, T., Kato, T. A., Kanba, S., Horikawa, H., Usami, M., Kodaira, M., Watanabe, K., Yoshikawa, T., Toyota, T., Yokoyama, S., Munesue, T., Kimura, R., Funabiki, Y., Kosaka, H., Jung, M., Kasai, K., Ikegame, T., Jinde, S., Numata, S., Kinoshita, M., Kato, T., Kakiuchi, C., Yamakawa, K., Suzuki, T., Hashimoto, N., Ishikawa, S., Yamagata, B., Nio, S., Murai, T., Son, S., Kunii, Y., Yabe, H., Inagaki, M., Goto, Y. I., Okumura, Y., Ito, T., Arioka, Y., Mori, D., and Ozaki, N. (2022) Cross-Disorder Analysis of Genic and Regulatory Copy Number Variations in Bipolar Disorder, Schizophrenia, and Autism Spectrum Disorder. Biol Psychiatry

22. Iijima, T., Wu, K., Witte, H., Hanno-Iijima, Y., Glatter, T., Richard, S., and Scheiffele, P. (2011) SAM68 regulates neuronal activity-dependent alternative splicing of neurexin-1. Cell 147, 1601–1614

23. Najib, S., Martin-Romero, C., Gonzalez-Yanes, C., and Sanchez-Margalet, V. (2005) Role of Sam68 as an adaptor protein in signal transduction. Cell Mol Life Sci 62, 36–43

24. Sanchez-Jimenez, F., and Sanchez-Margalet, V. (2013) Role of Sam68 in post- transcriptional gene regulation. Int J Mol Sci 14, 23402–23419

25. Berg, M. G., Singh, L. N., Younis, I., Liu, Q., Pinto, A. M., Kaida, D., Zhang, Z., Cho, S., Sherrill-Mix, S., Wan, L., and Dreyfuss, G. (2012) U1 snRNP determines mRNA length and regulates isoform expression. Cell 150, 53–64

26. Graf, E. R., Zhang, X., Jin, S. X., Linhoff, M. W., and Craig, A. M. (2004) Neurexins induce differentiation of GABA and glutamate postsynaptic specializations via neuroligins. Cell 119, 1013–1026

27. Green, E. K., Rees, E., Walters, J. T., Smith, K. G., Forty, L., Grozeva, D., Moran, J. L., Sklar, P., Ripke, S., Chambert, K. D., Genovese, G., McCarroll, S. A., Jones, I., Jones, L., Owen, M. J., O’Donovan, M. C., Craddock, N., and Kirov, G. (2016) Copy number variation in bipolar disorder. Mol Psychiatry 21, 89–93

28. Ishii, T., Ishikawa, M., Fujimori, K., Maeda, T., Kushima, I., Arioka, Y., Mori, D., Nakatake, Y., Yamagata, B., Nio, S., Kato, T. A., Yang, N., Wernig, M., Kanba, S., Mimura, M., Ozaki, N., and Okano, H. (2019) In Vitro Modeling of the Bipolar Disorder and Schizophrenia Using Patient-Derived Induced Pluripotent Stem Cells with Copy Number Variations of PCDH15 and RELN. eNeuro 6

29. Molumby, M. J., Anderson, R. M., Newbold, D. J., Koblesky, N. K., Garrett, A. M., Schreiner, D., Radley, J. J., and Weiner, J. A. (2017) gamma-Protocadherins Interact with Neuroligin-1 and Negatively Regulate Dendritic Spine Morphogenesis. Cell Rep 18, 2702–2714

30. Steffen, D. M., Ferri, S. L., Marcucci, C. G., Blocklinger, K. L., Molumby, M. J., Abel, T., and Weiner, J. A. (2021) The gamma-Protocadherins Interact Physically and Functionally with Neuroligin-2 to Negatively Regulate Inhibitory Synapse Density and Are Required for Normal Social Interaction. Mol Neurobiol 58, 2574–2589

31. Chih, B., Gollan, L., and Scheiffele, P. (2006) Alternative splicing controls selective trans-synaptic interactions of the neuroligin-neurexin complex. Neuron 51, 171–178

32. Sato, Y., Iijima, Y., Darwish, M., Sato, T., and Iijima, T. (2021) Distinct Expression of SLM2 Underlies Splicing-Dependent Trans-Synaptic Signaling of Neurexin Across GABAergic Neuron Subtypes. Neurochem Res

33. Hanawa, H., Kelly, P. F., Nathwani, A. C., Persons, D. A., Vandergriff, J. A., Hargrove, P., Vanin, E. F., and Nienhuis, A. W. (2002) Comparison of various envelope proteins for their ability to pseudotype lentiviral vectors and transduce primitive hematopoietic cells from human blood. Mol Ther 5, 242–251

34. Tufi, S., Lamoree, M., de Boer, J., and Leonards, P. (2015) Simultaneous analysis of multiple neurotransmitters by hydrophilic interaction liquid chromatography coupled to tandem mass spectrometry. J Chromatogr A 1395, 79–87

35. Hsiao, J. J., Potter, O. G., Chu, T. W., and Yin, H. (2018) Improved LC/MS Methods for the Analysis of Metal-Sensitive Analytes Using Medronic Acid as a Mobile Phase Additive. Anal Chem 90, 9457–9464

