## Supplementary file for "SAM68-regulated ALE selection of Pcdh15 maintains proper synapse development and function"

**Supplementary Figures and Tables**

**Figure S1: SAM68-regulated ALE selections**

Additional information related to Figure 1.

**Figure S2: U1 snRNP distinctly regulates SAM68-regulated ALE selections**

Additional information related to Figure 3.

**Figure S3: Generation of Pcdh15 Δex6 mutant by genome editing and its evaluation**

Additional information related to Figure 5

**Supplementary Tables**

**Table S1. List of PCR primer sets**

**Supplementary Figures and Tables**

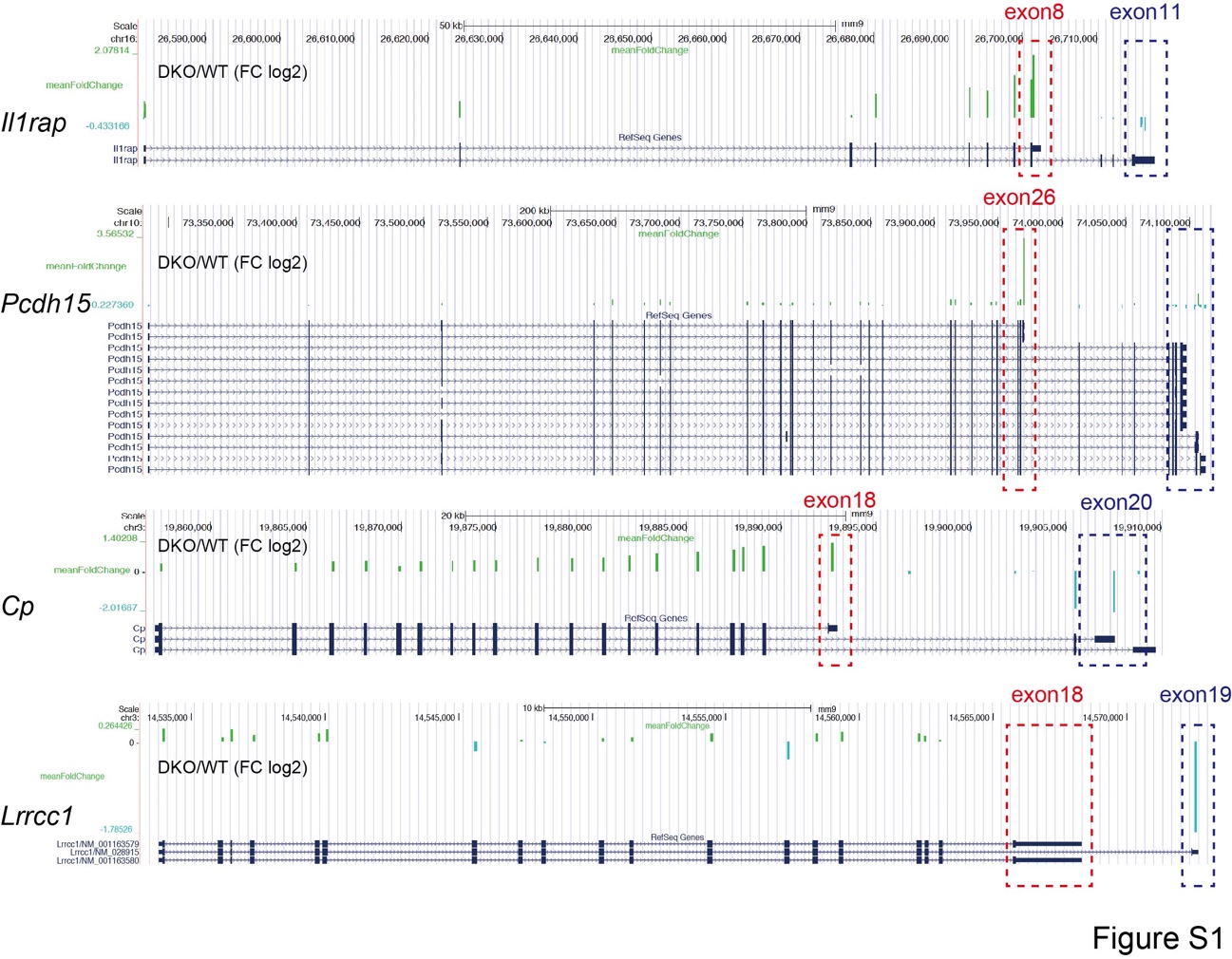

**Figure S1: SAM68-regulated ALE selections (Related to Figure 1)**

Atypical selection of the ALE of *Il1rap*, *Pcdh15*, *Cp*, and *Lrrcc1* genes in *Sam68*/*Slm1* ^DKO^ brains shown in exon array. Data of exon array based on the UCSC genome browser Mouse July 2007 (NCBI37/mm9) assembly.

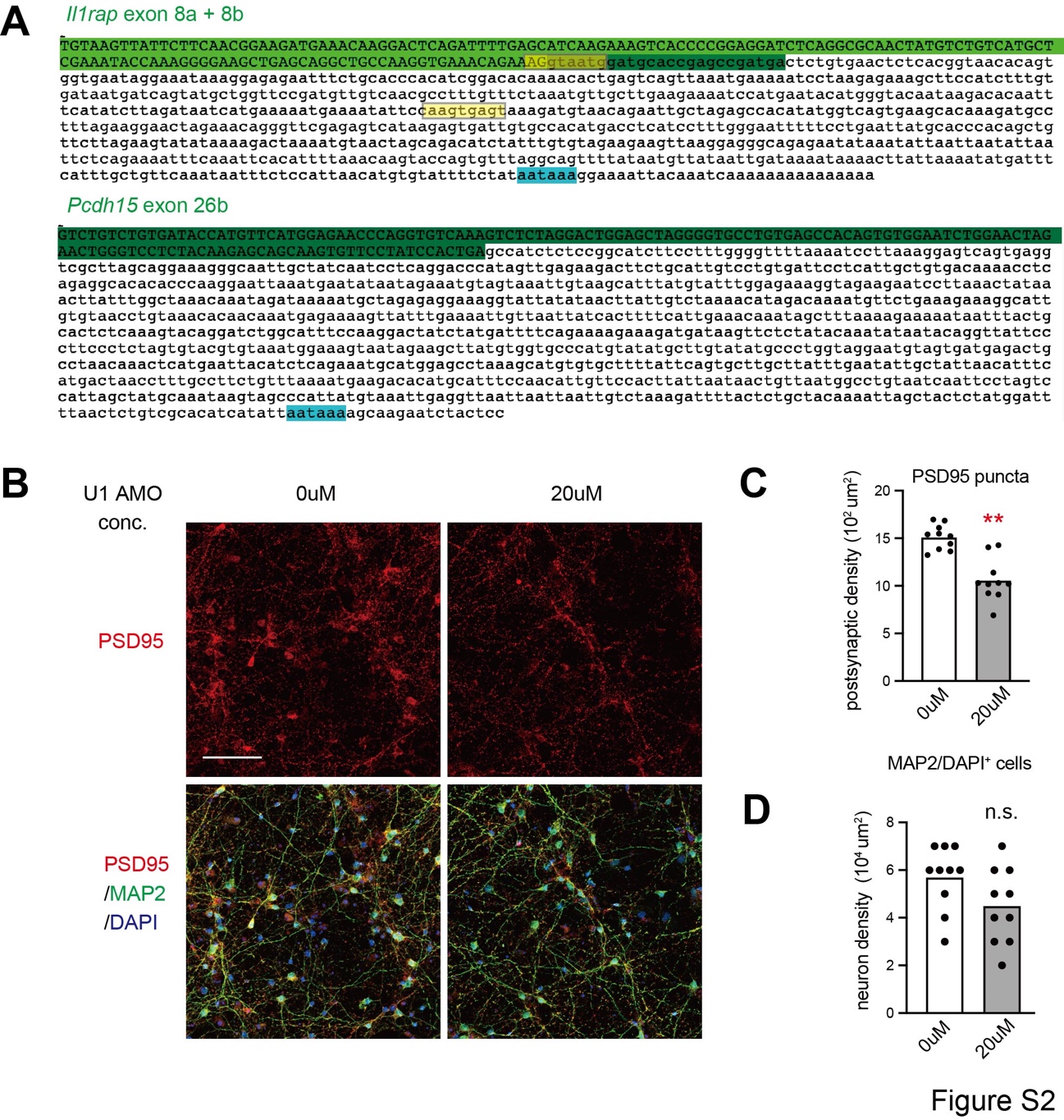

**Figure S2: U1 snRNP distinctly regulates SAM68-regulated ALE selections (Related to Figure 3)**

(A) The full-length cDNA sequences of *Il1rap* exon 8 and *Pcdh15* exon 26b. Green colors indicate the coding exon region. Blue colors show putative PAS sites on the 3’UTR. Yellow colors show putative U1 motifs.

(B-D) Immunostaining of cortical neuronal cultures at DIV10 electroporated with U1 AMOs (0 µM and 20 µM) with the neuronal markers PSD95 and MAP2. (B) Upper panels show PSD95-expressing puncta. Middle panels show overlay between PSD95 and MAP2 (C) The number of neurons (MAP2/DAPI^+^ cells). (n = 10 fields per group) (D) The density of PSD95-positive excitatory synapses electroporated with U1 AMOs (0 µM and 20 µM). (n = 10 fields per group). A student’s t-test was used.

Scale bars = 50 μm in (B).

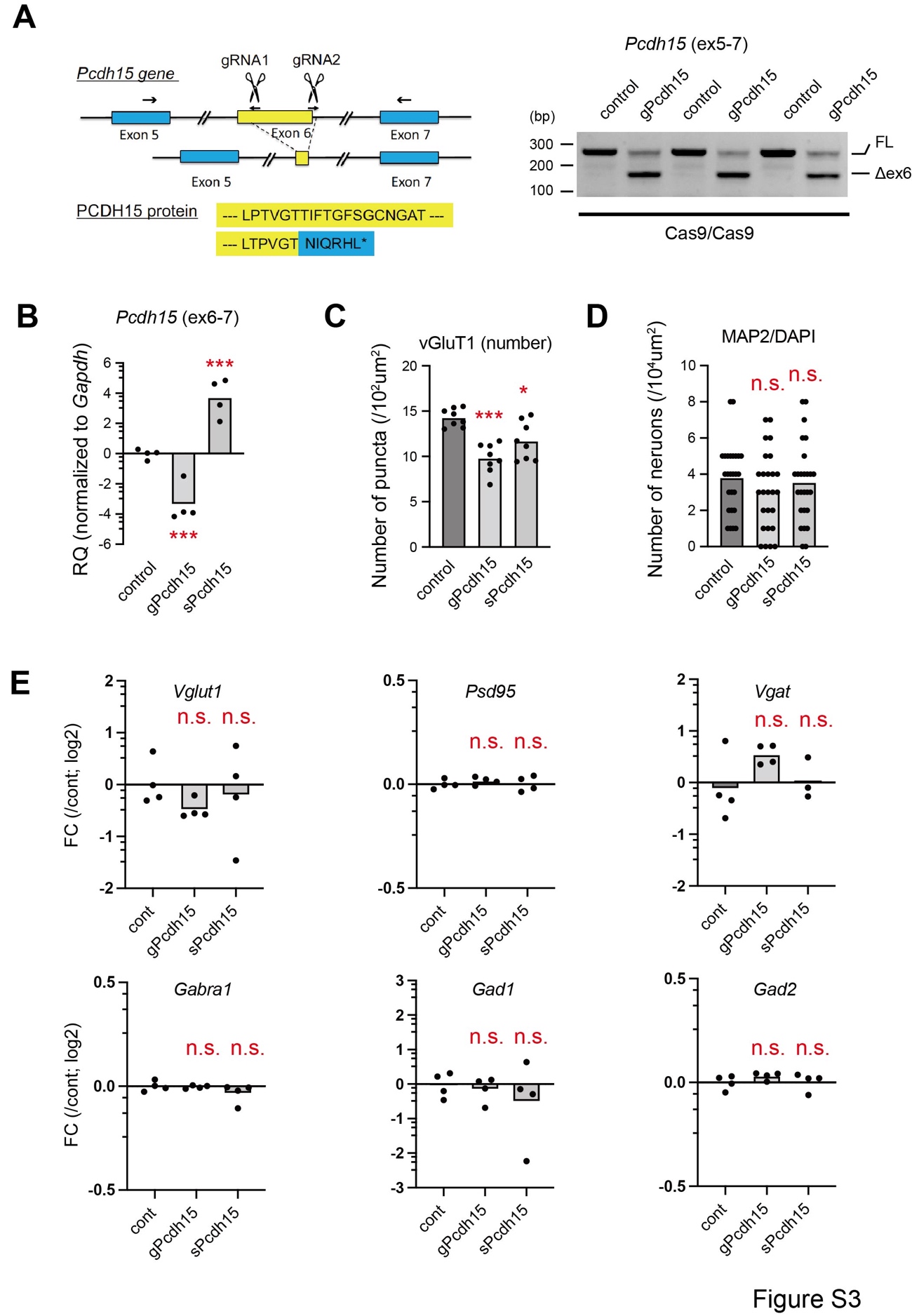

**Figure S3: Generation and characterization of *Pcdh15* Δex6 mutants using the CRISPR/Cas9 system (Related to Figure 5)**

(A) Left, schematic illustration showing the design of deleting *Pcdh15* gene locus with CRISPR/Cas9 system. The upper part shows the location of the designed gRNAs and subsequent effect on exon 6. Arrows show the design of primers used to confirm the deletions. The lower part shows the effect of exon 6 deletion on protein truncation. Right, semi-quantitative RT-PCR of *Pchd15* gene in neurons with Δex6 mutation (*gPcdh15*) and the controls.

(B) qPCR analysis of mRNA derived from neurons treated with lentiviral vectors carrying the control, *gPcdh15*, and *sPcdh15.* *Gapdh* was used as an internal control.

(C) The density of excitatory synapses treated with lentiviral vectors carrying the control, *gPcdh15*, and *sPcdh15*. n = 8 fields per group. A one-way ANOVA [F (2, 21) = 15.09, P <0.0001], followed by Turkey’s multiple comparisons test.

(D) The number of neurons (MAP2^+^/DAPI^+^ cells) counted in all the images used for the culture studies. n = 26 fields per group. One-way ANOVA [F (2, 75) = 0.82, p=0.44], followed by Turkey’s multiple comparison test. (E) qPCR analysis of mRNA of major synaptic markers derived from neurons treated with lentiviral vectors carrying the control, gPcdh15, and sPcdh15. Gapdh was used as an internal control. n = 4 cultures per group. A one-way ANOVA test was used.
