## Supplementary material for "SAM68-regulated ALE selection of Pcdh15 maintains proper synapse development and function": Supplemntary table 1

**RNA oligonucleotide sequences for guide RNAs**

| Forward primer  Reverse primer | Sequence |
| --- | --- |
| Pcdh15 gRNA1 S  Pcdh15 gRNA1 AS | 5'- ATC CGA TCC GGT AAG TAC TCA GGG -3'  5'- AAA CCC CTG AGT ACT TAC CGG ATC -3' |
| Pcdh15 gRNA2 S  Pcdh15 gRNA2 AS | 5'- ACC GCG AGA ACC CCG TGA ATA TCG -3'  5'- AAA CCG ATA TTC ACG GGG TTC TCG -3' |

**Oligonucleotide sequences of primer sets for semi-quantitative PCR**

| Forward primer  Reverse primer | Sequence | products (bp) |
| --- | --- | --- |
| Pcdh15 ex5-F  Pcdh15 ex7-R | 5'- TGT TGG GAC AGA TGA CAT CGC C -3'  5'- GAG AGC TGG CCC TGG AAG GG -3' | 250/158 |
| Gapdh-F  Gapdh-R | 5'- TGT TGC CAT CAA TGA CC -3'  5'- TCT CAT GGT TCA CAC CCA -3' | 342 |

**Oligonucleotide sequences of primer sets for RT-qPCR**

| Forward primer  Reverse primer | Sequence | products (bp) |
| --- | --- | --- |
| Pcdh15 ex25-F  Pcdh15 ex26a-R  Pcdh15 ex26b-R | 5’- CCG GGT ACA AGC AGA TTC TC -3’  5’- GGG TGA TCG TTT TCA TCC TG -3’  5’- TTG ACA CCT GGG TTC TCC AT -3’ | 108/101 |
| Pcdh15 total (ex25)-F  Pcdh15 total (ex25)-R | 5’- TGG ATT ACG AGA CAA GGA CCA -3’  5’- TTG AAG GGA CTC GGA GAT TG -3’ | 87 |
| Cp ex17-F  Cp ex17b-R  Cp ex19-R | 5’- GCT GGG ATG GCA ACT ACC TA -3’  5’- CAG TTG TGT GGC TTG GAT TTT -3’  5’- TTT TCC TGG CTA CTC CTT GG -3’ | 148/149 |
| Cp total-F  Cp total-R | 5’- GGT TCC TTC ACA AAC CGA AA -3’  5’- TGA ATG CTG AGA GGA TGC TG -3’ | 137 |
| Lrrcc1 ex18-F  Lrrcc1 ex19a-R  Lrrcc1 ex19b-R | 5’- GCG CAC CAA GCT GAA ATA AT -3’  5’- TTG CAT GTT TCG TCC AGA AG -3’  5’- GGG TGG TGT TTT TCA GTC TCA -3’ | 119/111 |
| Lrrcc1 total (ex18)-F  Lrrcc1 total (ex18)-R | 5’- TGC AAT GGA AAA GCT TCA GA -3’  5’- TCT GCT TCT CAT TTG CTA GCT G -3’ | 107 |
| Il1rap ex8-F  Il1rap ex9a-R  Il1rap ex9b-R | 5’- GCT GCC AAG GTG AAA CAG A -3’  5’- GGA CCA TCT CCA GCC AGT AA -3’  5’- GTG TTT TGT GTC CGA TGT GG -3’ | 130/127 |
| Il1rap total (ex3)-F  Il1rap total (ex3)-R | 5’- ACT ACA GCA CTG CCC ATT CC -3’  5’- CGG AAC CAG AGC ACA TCT TT -3’ | 136 |
| Sam68-F  Sam68-R | 5’- AAG AAC GCG TGC TGA TAC CT -3’  5’- GCA CCA GTC TCT TCC TGG AG -3’ | 109 |
| Slm1-F  Slm1-R | 5’- AAC GAG GAT GCC TAC GAC AG -3’  5’- TAG GGG TGC TCC CTG TAT CC -3’ | 110 |
| Pcdh15-6F  Pcdh15-7R2 | 5’- CAC GAT ATT CAC GGG GTT CT -3’  5’- GTC GTT GGA TGT CGG ATC TT -3’ | 115 |
